# Urocortin-3 neurons in the perifornical area mediate the impact of chronic stress on female infant-directed behavior

**DOI:** 10.1101/2022.02.19.481074

**Authors:** Brenda Abdelmesih, Robyn Anderson, Ilaria Carta, Anita E. Autry

**Author notes:** Correspondence should be addressed to Anita E. Autry.

## Abstract

Infant avoidance and aggression are promoted by activation of the Urocortin-3 expressing neurons of the perifornical area of hypothalamus (PeFA^Ucn3^) in male and female mice. PeFA^Ucn3^ neurons have been implicated in stress, and stress is known to reduce maternal behavior. We asked how chronic restraint stress (CRS) affects infant-directed behavior in virgin and lactating females and what role PeFA^Ucn3^ neurons play in this process. Here we show that infant-directed behavior increases activity in the PeFA^Ucn3^ neurons in virgin and lactating females. Chemogenetic inhibition of PeFA^Ucn3^ neurons facilitates pup retrieval in virgin females. CRS reduces pup retrieval in virgin females and increases activity of PeFA^Ucn3^ neurons but does not affect maternal behavior in mothers. Inhibition of PeFA^Ucn3^ neurons blocks stress-induced deficits in pup-directed behavior in virgin females. Together, these data illustrate the critical role for PeFA^Ucn3^ neuronal activity in mediating the impact of chronic stress on female infant-directed behavior.

**Significance statement:** While a large body of research has studied the impact of maternal stress on offspring, few studies have focused on the neural circuitry underlying reduced maternal behavior in stressed mothers. In this study, we examine the neural substrates involved in reduced infant-directed behavior caused by chronic stress. We find that perifornical area neurons expressing the neuropeptide urocortin-3 are critical mediators of the impact of stress on infant-directed behavior in females.

## Introduction

Many decades of research have focused on the neurobiology of maternal behavior, revealing common mechanisms and pathways involved in infant caregiving behavior across a variety of species (Numan & Insel, 2003; Numan, 2020). Studies have converged on the critical role of the medial preoptic area of hypothalamus in orchestrating the behavioral responses of mothers to their young in frogs, fish, birds, and rodents (Numan, 1974; Slawski & Buntin, 1995; Fischer *et al*., 2019; Maruska *et al*., 2020). Recently, it has been appreciated that these mechanisms may also be involved in paternal behaviors as well, suggesting a core circuitry that exists in both sexes to promote caregiving (O’Connell *et al*., 2012; Wu *et al*., 2014; Kohl *et al*., 2018). Moreover, it has been well-documented that neural plasticity mechanisms underlie the facilitation of infant care behavior, including alloparental care towards unrelated young, particularly in females (Numan & Insel, 2003).

In the absence of caregiving behavior, it is possible to observe neglect or even aggression toward infants by adults. Studies have identified circuit nodes in the brain, including the medial and posterior amygdala, the bed nucleus of stria terminalis, and the perifornical area of hypothalamus, that modulate expression of infant-directed neglect and aggression (Tsuneoka *et al*., 2015; Chen *et al*., 2019; Sato *et al*., 2020; Autry *et al*., 2021). We wondered if this anti-parental circuitry may be active in neglectful animals including virgin females or stressed virgin and lactating females. Clinical research clearly shows that stress is a critical risk factor for postpartum mental illnesses including postpartum depression or anxiety which affect up to 25% of women and 10% of men annually in the United States (Paulson & Bazemore, 2010; Wisner *et al*., 2013). However, there are few preclinical studies that examine the neurobiology underlying reduced parent-infant bonding or associated symptoms in animal models (Nephew & Bridges, 2011; Zoubovsky *et al*., 2020; Rosinger *et al*., 2021). Extant research has focused on the impact of maternal stress as a model of early life stress either pre- or postnatally on behavior outcomes in offspring, often with profound behavioral, physiological, and neurobiological impacts on young raised by stressed mothers (Cameron *et al*., 2005; Wang *et al*., 2011; Singh-Taylor *et al*., 2015; Delpech *et al*., 2016; Feifel *et al*., 2017; Kronman *et al*., 2021; Rincon-Cortes & Grace, 2021). During lactation, females are hyporesponsive to acute stress due to hormonal changes that impact the Hypothalamic-Pituitary-Adrenal (HPA) axis regulation (Walker *et al*., 2001; Brunton *et al*., 2008). HPA axis hypo-responsivity is thought to be protective of anxiety-related behavior and adult-pup interactions in lactating females (Miller *et al*., 2011; Medina *et al*., 2021). However, chronic stress has been documented to have a long-lasting impact on the regulation of the HPA axis, leading to reduced parenting, and how the underlying neurobiology is affected remains poorly understood (Carini *et al*., 2013; Murgatroyd & Nephew, 2013; Murgatroyd *et al*., 2015).

Ucn3 is a member of the corticotropin releasing factor (CRF) family of stress hormones and has the highest endogenous binding affinity for CRF receptor 2 (CRFR2). Previous studies of this group of neurons suggest that they are sensitive to stress and adrenalectomy (loss of stress hormones) (Jamieson *et al*., 2006). Overexpression of Ucn3 in the brain leads to increased anxiety- and depression-related behaviors and results in a blunted HPA response to stress (Neufeld-Cohen *et al*., 2012). Furthermore, overexpression of Ucn3 specifically in the PeFA is associated with enhanced anxiety-like behaviors in mice (Kuperman *et al*., 2010). Social discrimination abilities are altered in a sex-specific manner in total Ucn3 knockout mice (Deussing *et al*., 2010). Taken together, these studies suggest that PeFA Ucn3 cells mediate stress-induced behavioral changes.

Thus, we hypothesized that chronic stress would negatively affect infant-directed behavior in females and that this disruption is dependent on activation of perifornical area urocortin-3 expressing neurons. We set out to determine if anti-parental circuit components, specifically the urocortin-3 positive neurons in the perifornical area of hypothalamus were more active in naïve or stressed females, and if we could recover parental behavior by blocking activation of this anti-parental circuit node. We find that increased parental behavior is accompanied by decreased activity in perifornical area urocortin-3 expressing neurons and blocking activity in these cells enhances parental behavior in naïve females. Chronic stress reduces alloparental behavior in naïve females and this stress-induced behavioral effect is occluded by inhibition of perifornical area urocortin-3 cells. In stressed lactating females, parental behavior is preserved and perifornical area urocortin-3 cells are less activated in stress. Together, these data reveal a critical role for perifornical urocortin-3 neurons in the expression of alloparental behavior in female mice under both normal and pathological conditions.

## Materials and Methods

### Animals

Mice were maintained on a 12h:12h dark light cycle (10:30am-10:30 pm dark phase) with access to food and water ad libitum. All experiments were performed in accordance with NIH guidelines and approved by the Albert Einstein College of Medicine Institutional Animal Care and Use Committee (IACUC; protocol 20180110; 20180111; 00001386). C57BL/6J sexually naïve female and pregnant female (E14) mice were ordered from Jackson Laboratories (Bar Harbor, ME) aged at 6-8 weeks. Ucn3::Cre BAC transgenic line (STOCK Tg(Ucn3-cre) KF43Gsat/Mmcd 032078-UCD; obtained from laboratory of Catherine Dulac, Harvard University) were genotyped at weaning (3 weeks of age) and used in experiments at age 2-5 months. Animals received from Jackson Laboratories habituated to our facility for 7 days prior to behavioral testing.

### Corticosterone Measure

Trunk blood samples were taken at the time of sacrifice and blood serum was isolated from blood samples by centrifugation. A high-sensitivity corticosterone (CORT) enzyme immunoassay (EIA) was used and analyzed according to manufacturer’s instructions (Immunodiagnostic Systems Ltd, Fountain Hills, AZ, USA) as previously described (Autry *et al*., 2009). Briefly, percent binding (B/Bo%) of each calibrator, control and sample was calculated by dividing the mean absorbance over the mean absorbance for ‘0’ calibrator and multiplied by 100. A calibration curve was used to plot B/Bo% on the ordinate against concentration of corticosterone. A 4pl curve fit was applied.

### Chronic restraint stress model

Virgin and lactating female mice were used. Lactating females were restrained starting from approximately postpartum (PP) day 2. Both stressed and unstressed mice were brought to a test room under dim red light during their dark cycle. All animals were weighed, and females were either placed back into their home cage or placed into a 50 mL conical tube for one hour. Humidity and temperature in the test room was recorded each day. On the last day of stress, females remained in the test room for 1-2 hours before being exposed to a foreign-born pup (see Parental Behavior).

Animals injected with AAV1/DIO-hM4Di (see chemogenetics) recovered from surgery at least 1 week before the start of restraint. For the stressed virgin female group, mice were excluded from the stressed group (n=4 females) based on open field test behavior that was indistinguishable from control. For the groups that received AAV1/DIO-hM4Di injections, females were excluded if they did not show adequate recombination of the DREADD construct (n=2, virgin females Figure 2; n=0 stressed virgin females Figure 6).

### Intruder stress model

Lactating females were stressed starting from approximately postpartum (PP) day 2. Both stressed and unstressed mice were brought to a test room under dim red light during their dark cycle. All animals were weighed, and females were placed back into their home cage and either placed back on the housing rack or had an intact adult male intruder (C57, ∼2 months old) introduced into their home cage for 10 minutes as described in previous studies (Carini *et al*., 2013; Murgatroyd *et al*., 2016). In the event the male was too aggressive toward pups, the intruder stress period was curtailed. Humidity and temperature in the test room was recorded each day. On the last day of stress, females remained in the test room for 1-2 hours before being exposed to a foreign-born pup (see Parental Behavior).

### Behavior assays

Mice were individually housed for at least 1 week prior to testing. Experiments were conducted during the dark phase under dim red light. Tests were recorded by Fly Capture cameras (Point Grey, Richmond, BC, Canada) and behaviors were scored by an observer blind to experimental condition using Observer XT13 Software or Ethovision XT 13 (Noldus Information Technology, Leesburg, VA, USA). Animals were tested for a single behavior per session with at least 24 hours between sessions.

### Parental behavior

Parental behavior tests were conducted in the mouse’s home cage as previously described (Wu *et al*., 2014). Mice were habituated to the testing environment for 10 minutes. One to two C57BL6/J pups 1-4 days old were presented in the cage in the opposite corner to the nest. Test sessions started either at pup introduction or pup approach (female first touches the pup with its snout) and lasted for 10-15 minutes. If the mouse became aggressive by biting and wounding the pup, the session was immediately halted, and the pup was euthanized. The following behaviors were quantified: latency to retrieve, pup investigation (sniffing, close contact with snout), grooming (handling with forepaws and licking), nest building, time spent in the nest, crouching, latency to attack (latency to bite and wound), aggression (roughly handling, aggressively grooming, aggressive carrying with no retrieval), and tail rattling. A ‘parenting behavior’ index was calculated as the sum of duration of grooming, nest building, time spent in the nest, and crouching.

### Open field

Mice were assessed for activity in a 45cm x 45 cm open field at 40 lux for 5 min as previously described (Autry *et al*., 2009). Center was considered 15 cm x 15 cm and borders were 5 cm around the perimeter of the box. Time and frequency in center and borderas well as distance and velocity were calculated using Ethovision XT13. Behavioral ethograms were made in Matlab using custom code.

### Fluorescence in situ hybridization

Fluorescence in situ hybridization (FISH) was performed as recommended by ACD Bio (Newark, CA, USA) using V1 RNAscope reagents. Briefly, fresh brain tissue was collected from animals housed in their home cages or 35 min after the start of the behavior tests for immediate early gene (*Fos)* studies. Brains were embedded in OCT (Tissue-Tek) and frozen with dry ice. 25μm cryosections were used for mRNA in situ. Adjacent sections from each brain were collected over replicate slides to stain with multiple probes. Protease 3 was used to digest tissue. Fos (Cat No. 316921), Ucn3 (Cat No. 464861), and Crh (Cat No. 316091) probes were used as per manufacturer’s instructions. Slides were mounted using Prolong Gold with DAPI. Zeiss Axioscan was used to image DAPI, Alexa 488, Atto-550, and Alexa 647 at 20X magnification.

### Immunostaining and histology

To visualize c-Fos protein in combination with AAV-hM4Di, perfused tissue was sliced on a freezing microtome at 30 μm, and every third section throughout the PeFA was stained. Sections were rinsed with 0.1% PBS with Triton (PBST), blocked with 5% donkey serum diluted in PBST (blocking solution) for 1 hour at room temperature. Primary antibody chicken anti-mCherry (Millipore AB3566481) and rabbit anti c-Fos (Cell Signaling 2250S) were diluted at 1:1000 in blocking solution and sections were incubated overnight at 4°C. After rinsing with PBST, secondary anti-chicken-A594 (Sigma CF594) and anti-rabbit-A647 (Life Technologies A31573) were applied at 1:200 and 1:1000 dilutions, respectively, in blocking solution and incubated overnight at 4°C. Sections were rinsed in PBS, mounted to Superfrost Plus slides, coverslipped with Prolong Gold containing DAPI, and imaged on the Zeiss Axioscan as described previously.

### Chemogenetics

Ucn3::Cre virgin female mice (or Cre negative littermates as controls) 8-20 weeks old were used for these experiments. We stereotaxically injected ∼225 nL of conditional inhibitory designer receptor exclusively activated by designer drug (DREADD) virus bilaterally into the PeFA (AP −0.6mm, ±ML 0.3mm, DV −4.2mm). For the naïve virgin female DREADD experiment, we used a custom prep from UNC Vector Core (AAV1-hSyn-DIO-HM4D(Gi)-mCherry; Chapel Hill, North Carolina, USA) and for the stressed virgin female DREADD experiment we used a custom prep from Vector Builder (AAV1-hSyn-FLEX-HM4D(Gi)-mCherry; Chicago, Illinois, USA). Animals used for stress study recovered from surgery for 1 week before the start of restraint and around two-three weeks before behavioral testing.

Cre-positive and Cre-negative females were administered intraperitoneally (i.p) with either 1x PBS (vehicle) or 0.3mg/kg clozapine-n-oxide (CNO) dissolved in 1x PBS and habituated to the testing environment for two-three hours prior to pup assay. Females were presented one to two C57BL6/J pups in the corner of their home-cage opposite the nest and parental behaviors were recorded for 10-15 minutes.

Four control animals used in the stress study were administered either CNO or saline i.p and 2 hours later were exposed to a pup. Animals were then perfused 90 minutes later to stain for c-Fos protein expression. (See Immunostaining and histology)

### Data analysis and Statistics

Data was analyzed by Graphpad Prism 9.0 or Matlab scripts. For colocalization experiments, we used Fisher’s exact test to compare the total number of fos+/marker+ positive cells to the total number of fos-/marker+ positive cell populations across all mice and expressed the data as percentages from each individual mouse. Pup retrieval percentages are analyzed by Kolmogorov-Smirnoff (2 groups) or Friedman test (3 groups). For experiments comparing one manipulation to control (i.e., stress or neuronal inactivation), we used t-test for parameters with normally distributed data and Mann-Whitney test for non-normally distributed data. To compare one manipulation and control across several sessions, we used one-way repeated measures ANOVA tests followed by post-hoc correction. In experiments with comparison of two manipulations in several sessions (stress and neuronal inactivation), we used two-way repeated measures ANOVA followed by post-hoc correction. P values reported as follows: <0.05 *, ** P<0.01, *** P<0.001, **** P<0.0001. All data are expressed as mean ± SEM.

### Image analysis

Images were exported from Zen Blue software and cells were manually counted for colocalization using FIJI Cell Counter. Graphpad Prism 9 was used to plot graphs and perform statistics.

### Fiber Photometry

Ucn3-cre animals were injected with a cocktail of 150 nL of AAV-syn-jGCaMP7f-WPRE (Addgene 104488-AAV9) and 225 nL of AAV-hSyn-DIO-hM4D(Gi)-mCherry into PeFA (ML: 0.3mm; AP −0.6mm; DV 4.2mm). In the same surgery, a 200µM fiber optic cannula was implanted (ML: 0.3mm; AP −0.6mm; DV 4.2mm). Animals recovered for at least 3 weeks before behavioral experiments. Animals were brought up to the test room in dim red light and injected with either vehicle (session 1) or 0.3mg/kg CNO (session 2) i.p. Two hours later, a fiber optic patch cable (Doric) was attached to the cannula and adjusted to attachment for 10 minutes before recording. Using a multi-channel fiber photometry system (Neurophotometrics LTD), a 470 nm LED and 415 nm LED (isosbestic control) alternatively illuminated at 60µW via a 20X objective and fluorescence emission was collected using a CMOS camera sensor. After 1-2 minutes of recording, animals underwent 6 tail suspensions for approximately 5 seconds per suspension. Data were acquired using the open-source software Bonsai.

Photometry data was analyzed using custom MATLAB code. To correct for photobleaching and motion artifact, we used normalization similarly described by Hvratin et al 2020 (Hrvatin *et al*., 2020). In short, the isosbestic signal was fit with a biexponential and then linearly scaled to fit signal emitted by GCaMP. GCaMP signal was then divided by the scaled fit ΔF/F. Tail suspension events were aligned to normalized photometric signal and peri-events were taken from 5 sec before tail suspension (“pre”) to 5 sec after (“post”). The pre-event baseline was used to calculate the z score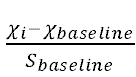. The mean ΔF/F of each pre- and post-tail suspension event was taken and averaged across animals per group (vehicle vs CNO) and compared using a paired t test. Area under curve was calculated with the mean pre- and post-tail suspension event using the MATLAB built-in function “trapz”. Standard error mean is plotted with the average z-score.

### Code availability

Custom Matlab code for ethogram generation and analysis of photometry data is available upon request.

## Results

To identify the activation levels of Urocortin-3 in the rostral perifornical area of the hypothalamus, we exposed C57 virgin females as well as lactating females (postpartum day 2) to either a foreign pup (P0-P4) or ∼25 mg of fresh bedding (control) in their home cage. Animals were subsequently sacrificed 30 minutes after exposure (Figure 1A). To control for number of pups as well and foreign pup discrimination (Ostermeyer & Elwood, 1983; Mogi *et al*., 2017), we utilized two groups of lactating females that either had their litter removed 10 minutes prior to foreign pup introduction or kept their litter (Figure 1A). Visualization of immediate early gene Fos and Ucn3 in PeFA revealed increases in PeFA^Ucn3^ cell activation in virgin females exposed to a pup compared to controls (Figure 1B, C; Supplemental Figure 1-1). However, in lactating females, Fos levels in PeFA^Ucn3^ decrease with pup exposure if litter has been removed but increase if litter is present. These results suggest that PeFA^Ucn3^ cells respond to pup exposure similarly in virgin females and lactating females that keep their litter, while we observe opposite impact on PeFA^Ucn3^ cell activity in mothers when her litter is removed.

**Figure 1.**
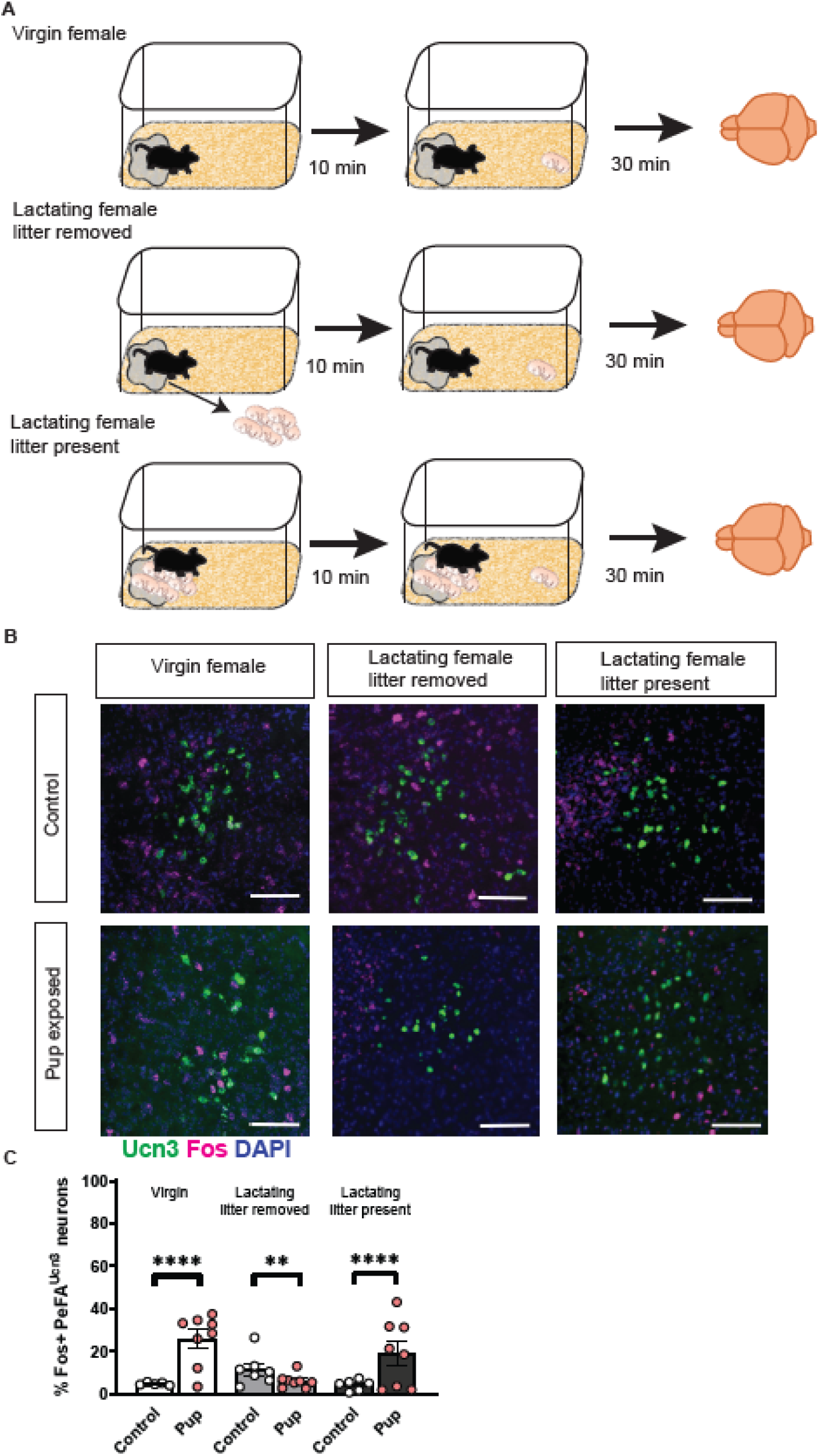
PeFA Urocortin-3 neuronal activation levels in response to foreign pups depends on physiological context. **(A)** Schematic of behavioral paradigm. C57 virgin females were exposed to a newborn pup and sacrificed 30 minutes after pup exposure or addition of fresh bedding into home cage (control). Lactating females either had a litter removed or litter intact and exposed to a foreign-born pup or fresh bedding. **(B)** Rostral perifornical area cells containing *Urocortin-3* and *Fos* RNA were counted for colocalization in each group. (scale bar 100 µm**) (C)** Quantification of percentage of Ucn3+ cells colocalized with Fos across groups. Fisher exact test reveals that pup exposure increases PeFA^Ucn3^ activation in virgin females (Control n=605 N=5; Pup exposed n=655 N=8; ****p=<0.0001), reduced PeFA^Ucn3^ activation in lactating females with litter removed (Control n=622 N=7; pup exposed n=828 N=8; **p=0.0044), and increased activation in lactating females that did not have litter removal compared to control bedding exposure (Control n=816 N=6; pup exposed n=988 N=8 ****p<0.0001).

Because virgin females are not as parental as lactating females (Lonstein & De Vries, 2000; Kuroda *et al*., 2011; Marlin *et al*., 2015; Carcea *et al*., 2021) and PeFA^Ucn3^ neurons are activated by infanticide in females (Autry *et al*., 2021), we wanted to test if suppression of PeFA^Ucn3^ neuronal activity could enhance alloparental behavior in virgin females. To accomplish this, we used a conditional viral strategy to express the inhibitory designer receptor exclusively activated by designer drug (DREADD hM4Di) in Ucn3::Cre positive and Cre negative animals in the PeFA of virgin females naïve to pups. Two-three weeks after viral injection, both groups of animals were administered CNO (0.3 mg/kg intraperitoneally) and 2-3 hours later, exposed to two pups for fifteen minutes in their home cage (Figure 2A; Supplemental Figure 2-1). We confirmed viral recombination to include females in subsequent behavioral analyses (Figure 2B). Cre+ females retrieved more pups in a shorter amount of time relative to Cre-females (Figure 2C & D). However, there was no difference in latency to retrieve the 2^nd^ pup (Figure 2F). Furthermore, Cre+ animals spent more time in the nest with pups and started nest-building earlier compared to Cre-females (Figure 2K &N). While other behaviors were not improved (Figure 2 G-J, L, M,O-P), suppression of PeFA^Ucn3^ neurons improved certain aspects of alloparenting behavior (Figure 2Q), particularly pup retrieval and time spent in the nest with the pups.

**Figure 2.**
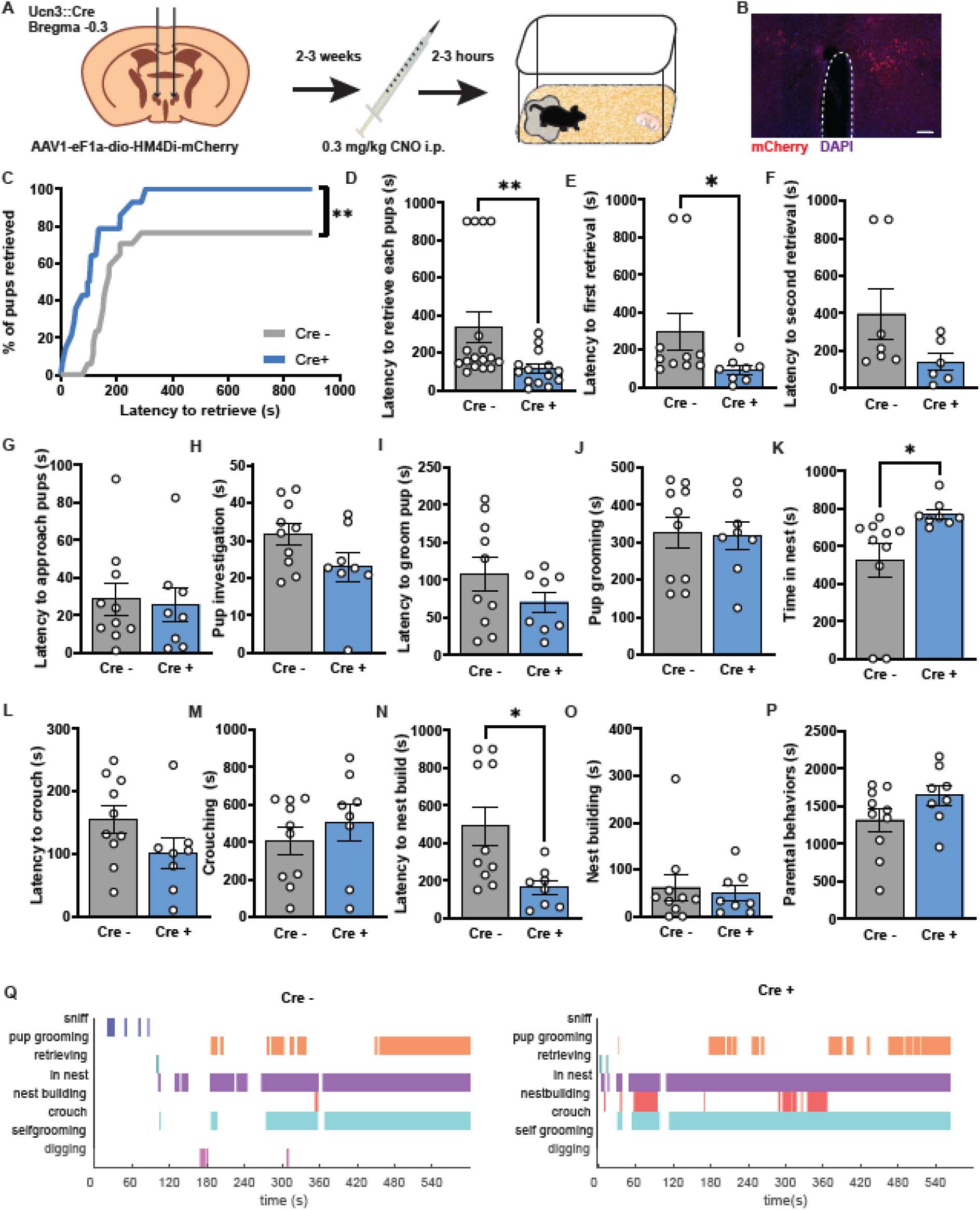
Inhibition of PeFA^Ucn3^ neurons enhances alloparental behavior in virgin female mice. **(A)** Schematic of viral injection strategy and behavior timeline (n=10 Cre-females; n=8 Cre+ females). **(B)** Representative image of mCherry reporter expression in Ucn3::Cre+ female injected with inhibitory DREADD virus (magenta: mCherry; blue: DAPI; scale bar 100 µm). **(C)** Percentage of pups retrieved by Cre+ females is significantly increased compared to Cre-females (Kolmogorov-Smirnov test p=0.0048). **(D)** Latency to retrieve pups is significantly faster in Cre+ females compared to Cre-females (Two-tailed Mann-Whitney test; p=0.0033), as well as **(E)** latency to retrieve the first pup (Two-tailed Mann-Whiney test; p=0.0259), but there was no difference between groups in **(F)** latency to retrieve the second pup. **(G-H)** Latency to approach pups was not significantly different, but time spent investigating pups trended lower in Cre+ animals (Unpaired t test; p=0.0854). **(I-J)** We observed no significant difference in latency to pup groom or in time spent pup grooming in Cre+ animals compared to Cre-females. **(K)** Time spent in nest with pups significantly increased in Cre+ animals (Unpaired t test; p=0.0289). **(L-M)** Latency to crouch trended lower in Cre+ animals (Two-tailed Mann-Whiney test; p=0.1011), but we did not observe a significant difference in time spent crouching. **(N-O)** Latency to nest build was significantly faster in Cre+ animals (Two-tailed Mann-Whiney test; p=0.0152) but time spent nest building was not significantly different between groups. **(P)** Cumulative time spent parenting was unchanged between groups **(Q)** Representative behavior trace of a Cre-animal (left) and a Cre+ animal (right) during pup assay (time 0 is when pup was added to home cage).

To understand the impact of chronic stress on alloparental behavior in virgin females, we employed a chronic restraint stress paradigm in which females were placed into a 50 mL Falcon conical tube for 1 hour a day for 20 days (Figure 3A). Stressed females weighed significantly less than control females (Figure 3B). On day 19, females were tested for exploratory behavior in a 5-minute open field task (Figure 3C-F). Stressed females spent less time in the center of the field compared to control females (Figure 3C). On the last day of stress, day 20, females were exposed to a newborn pup (P0-4) for 15 minutes and their alloparental behavior was recorded and analyzed (Figure 3G-R). Only 2 out of 9, or 22%, of stressed females retrieved pups compared to control females (5 of 9, or 55% retrieved) (Figure 3G). Other than retrieval latency, virgin stressed females did not show any significant changes in other measures of alloparental behavior. Altogether, we find that chronic restraint stress significantly reduces pup retrieval in virgin females.

**Figure 3.**
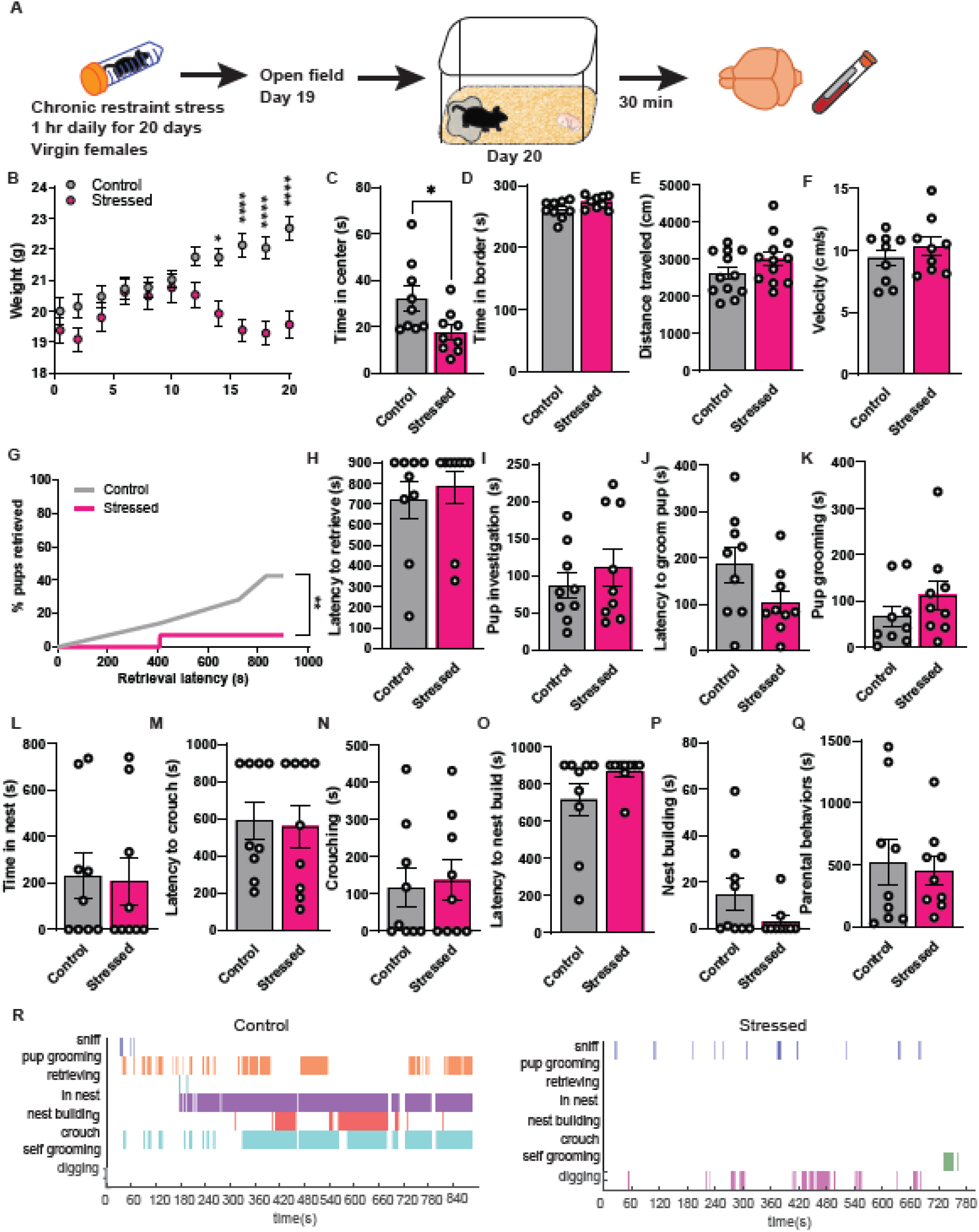
Chronic restraint stress dampens alloparental behavior in virgin female mice. **(A)** Schematic of timeline using restraint stress paradigm and behavioral testing in virgin females (n=9 control; n=9 stress). **(B)** Weights taken from each group during the period of chronic restraint stress. Stressed females have significant difference in weight compared to controls (Two-way repeated measures ANOVA; main effect of interaction of stress x time F_(10,220)_=22.92 p<0.0001; main effect of time F_(10, 220)_ = 20.35; Bonferroni’s multiple comparisons test) **(C-F)** Time spent in center of open field was significantly reduced for stressed females compared to controls (C) (unpaired t test; p=0.0289) but no other parameter in open field was changed. **(G)** Chronic restraint stress significantly decreased cumulative pup retrieval in females (Kolmogorov-Smirnov test p=0.0059).**(H-Q)** Chronic restraint stress did not significantly change other parenting measures such as retrieval latency, or time spent pup grooming, time in nest, crouching, and nest building. **(R)** Representative behavior trace for a control female (left) and stressed female (right) during pup assay. (*p=0.05; **p=0.01; ***p=0.001; ****p=0.0001).

Next, to understand the impact of chronic stress on maternal behavior in lactating females, we utilized the same chronic restraint paradigm in females from postpartum day 2-18 (Figure 4A), before weaning age for pups. Like stressed virgin females, stressed lactating females weighed significantly less than control females (Figure 4B), indicating that chronic restraint induced physiological changes. On day 16 of chronic restraint, females were tested for anxiety-related behavior in a 5-minute open field task (Figure 4C-F). Surprisingly, stressed females spent more time in the center of the field and less time in the borders (Figure 4C, D).

**Figure 4.**
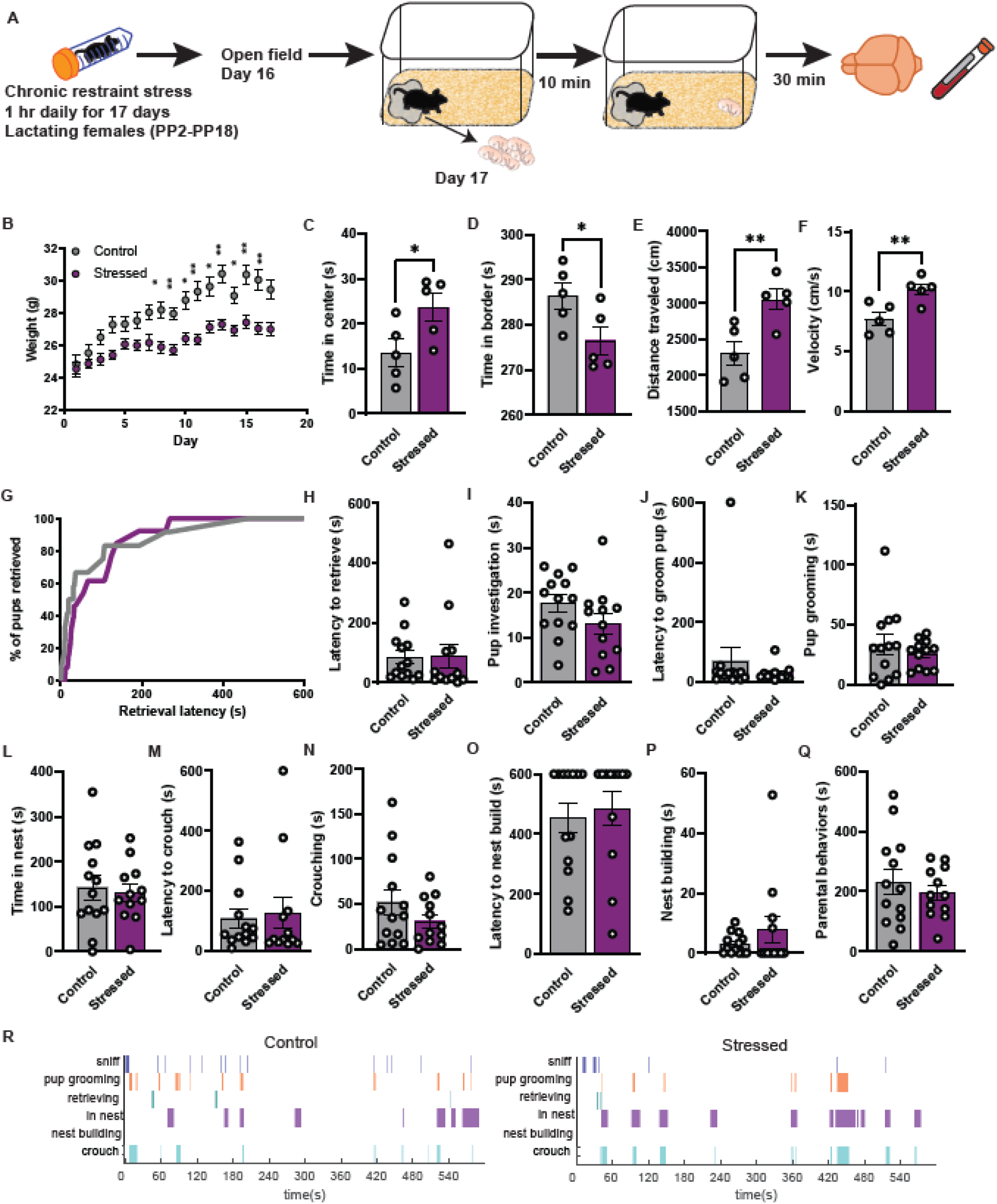
Chronic restraint stress does not induce changes in parental behavior in lactating females. **(A)** Schematic of timeline using restraint paradigm and behavioral testing in lactating females (Control n=12; Stressed n=13). **(B)** Weights taken from each group during the period of chronic restraint stress from postpartum day 1 to 18. Stressed females have a significant difference in weight compared to controls (Two-way repeated measures ANOVA; main effect of interaction of stress x time F_(17,368)_=8.286 p<0.0001; main effect of time F_(7.579,174.3)_=75.28 p<0.0001; Sidak’s multiple comparisons). **(C-F)** Time spent in center of open field was significantly increased for stressed females compared to controls (unpaired t test; p=0.0459) and time spent in borders decreased (unpaired t test; p=0.0459) accompanied by increased distance traveled (unpaired t test; p=0.0089) and velocity (unpaired t test; p=0.0089) **(G-Q)** Chronic restraint stress did not significantly change any parenting measures as retrieval, pup grooming, time in nest, crouching, and nest building compared to females that did not receive stress. **(R)** Representative behavior trace for a control female (left) and stressed female (right) during pup assay.

Stressed females also showed an increase in velocity and distance traveled relative to control females (Figure 4E, F). On the last day of stress, day 17, females had their litters removed and 10 minutes later we introduced a foreign-born pup to their home cage (Figure 4G-R). All females retrieved pups before the end of the 10 minutes session (Figure 4G, H) and there was no difference in latency to retrieve between groups. Stressed mothers showed similar levels of parenting toward pups as control mothers. We also attempted to use an intruder stress paradigm that has previously been reported to impact parental behavior in lactating females (Carini *et al*., 2013; Murgatroyd *et al*., 2016). We did not see any weight changes or parenting measures (Supplemental Figure 4-1).

Next, we investigated molecular and physiological impacts of chronic restraint stress in virgin or lactating females. In situ hybridization revealed increases in PeFA^Ucn3^/Fos colocalization in stressed virgin females (Figure 5A, B; Supplemental Figure 5-1). In control virgin females, percentage of PeFA^Ucn3^/Fos colocalization was negatively correlated with time spent parenting, indicating that activation of PeFA^Ucn3^ may reduce alloparental behaviors (Figure 5E). Because activation of corticotropin releasing factor (CRF) cells in the paraventricular hypothalamus (PVH) is postulated to disrupt maternal behavior and is critical for physiological stress responses (Herman & Tasker, 2016; Klampfl & Bosch, 2019), we also quantified PVH^CRF^/Fos colocalization (Figure 5C,F). We found that PVH^CRH^/Fos levels were significantly reduced in both stressed virgin females and lactating females, and like PeFA^Ucn3^, PVH^CRF^ neuronal activation is negatively correlated with parental behaviors (Figure 5F). Chronic restraint stress did not affect circulating CORT levels in virgin females (Figure 5D). In mothers, chronic stress led to a decrease in PeFA^Ucn3^/Fos levels compared to control lactating females, opposite to the effect we observed in virgin females (Figure 5G, H). Like virgin females, however, PVH^CRF^ cell activation was significantly decreased in chronically stressed lactating females (Figure 5G, I; Supplemental Figure 5-1), consistent with previous literature (Girotti *et al*., 2006; Radley & Sawchenko, 2015; Matovic *et al*., 2020). We observed a similar negative trend for correlation of PeFA^Ucn3^ neuronal activation and parental behaviors in control lactating females that we observed in virgin females, but the trend for PVH^CRF^ cell activation is positively correlated in lactating females (Figure 5 K, L). We observed that CORT levels were significantly decreased in stressed lactating females (Figure 5J), suggesting adaptive habituation to the repeated stress. In our intruder stress experiment, we did not observe molecular changes in PeFA^Ucn3^ activation or in circulating corticosterone levels, consistent with no changes in weight or parental behavior (Supplemental Figure 5-2). Altogether, chronic restraint stress induces differential PeFA^Ucn3^ activation patterns in virgin and lactating females in response to pups, while chronic stress reduces PVH^CRF^ neuronal activation in both virgins and mothers.

**Figure 5.**
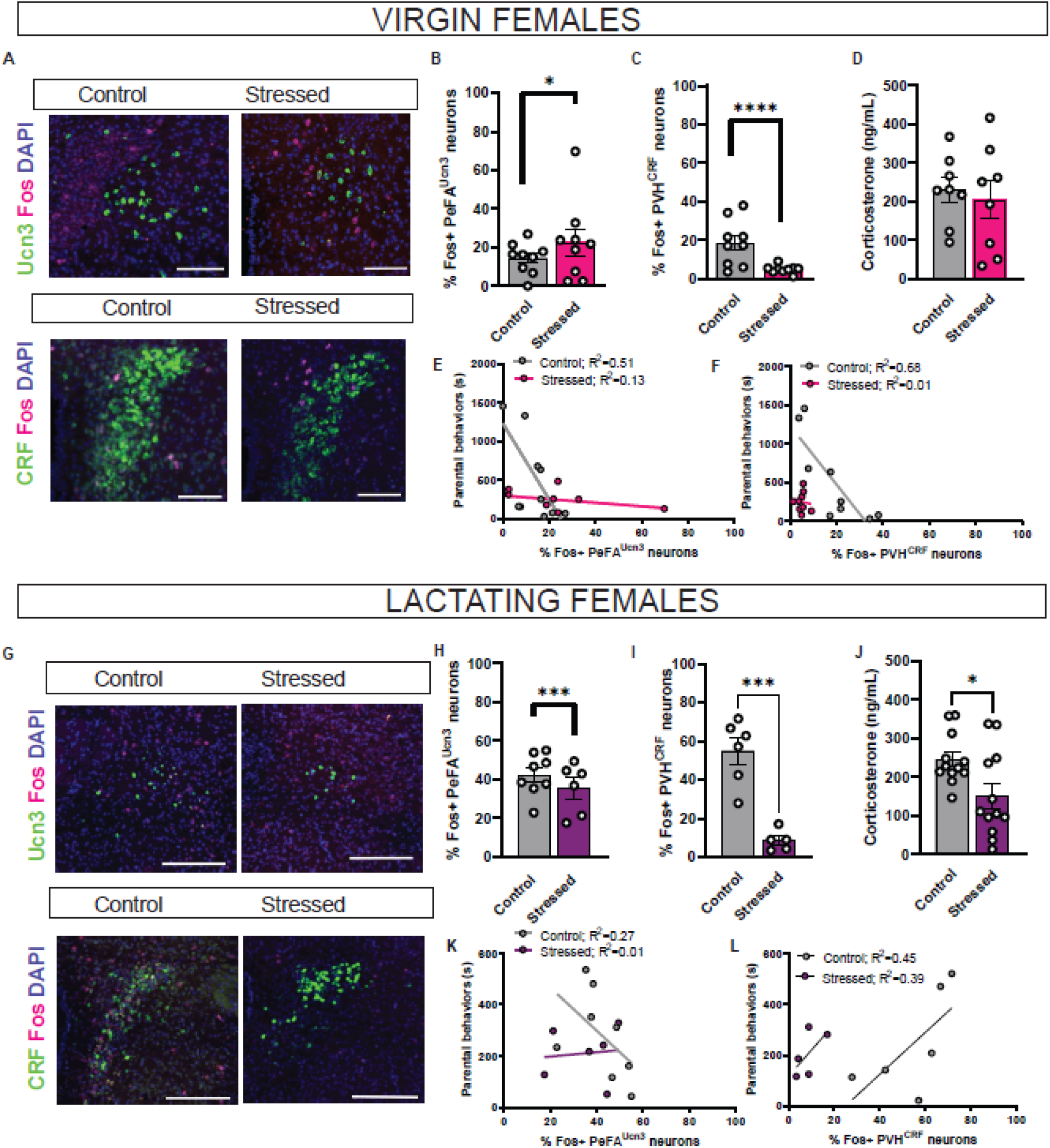
Chronic restraint leads to contrasting molecular changes between virgin and lactating females in parenting. **(A)** Rostral perifornical area cells containing *Ucn3, CRF,* and *Fos* RNA were counted for colocalization in each virgin female group. (scale bar 100 µm). **(B)** Colocalization of *Ucn3* and *Fos* in perifornical area reveals increased of activation of PeFA^Ucn3^ neurons in in stressed virgin females compared to control virgin females (Fisher exact test: Control n=490 N=9; Stressed n=472 N=9; p=0.038). **(C)** Colocalization of *CRF* and *Fos* in paraventricular nucleus of the hypothalamus reveals decreased activation of PVH^CRF^ neurons in stressed virgin females compared to control virgin females (Fisher exact test: Control n=1305 N=9; Stressed n=1272 N=9; p<0.0001). **(D)** Serum corticosterone levels were measured using ELISA in virgin females. Chronic restraint stress did not contribute to altered corticosterone in virgin females. **(E)** Plotting time spent parenting against PeFA^Ucn3^ activation levels shows marked negative correlation between parenting and PeFA^Ucn3^ neuron activation in controls but this relationship is abolished in stressed virgin females (Control R^2^=0.51 Stressed R^2^=0.13; difference in slope F= 10.73 DFn=1, DFd=14; p=0.0055). **(F)** PVH^CRF^ activation levels in negatively correlated with time spent parenting in control virgin females but this relationship is indiscernible in stressed virgin females (Control R^2^=0.68 Stressed R^2^=0.01). **(G)** Rostral perifornical area cells containing *Ucn3, CRF,* and *Fos* RNA were counted for colocalization in each lactating female group (scale bar 100 µm). **(H)** PeFA^Ucn3^ neuron activation levels are reduced in stressed lactating females (Fisher exact test: Control n=338 N=8; Stressed n=360 N=6; p=0.0005). **(I)** Activation levels of CRF cells in the paraventricular nucleus of the hypothalamus are significantly decreased in stressed lactating females (Control n=919 N=6; Stressed n=2268 N=5; p<0.0001). **(J)** Chronic restraint stress significantly reduced corticosterone in lactating females (unpaired t test; p=0.0198). **(K)** Linear regression analysis reveals a trend towards negative correlation between time spent parenting and PeFA^Ucn3^ activation levels in control lactating females but not in the stressed group (Control R^2^=0.27 Stressed R^2^=0.01). **(L)** PVH^CRF^ activation is positively correlated with time spent parenting (Control R^2^=0.45 Stressed R^2^=0.39).

Because our chronic restraint stress paradigm dampened alloparental behavior in virgin females and increased PeFA^Ucn3^ neuronal activation, we sought to ameliorate deficits in parenting by inhibiting PeFA^Ucn3^ neurons during pup exposure. To accomplish this, we injected virgin Ucn3::Cre positive and negative females with AAV1-eF1a-DIO-hM4Di-mCherry and then started restraint stress 1 week after recovery from surgery. After 16 days of chronic restraint stress, we injected either vehicle or CNO on 2 consecutive days after the last day of stress.

After several days we then employed randomized CNO/Vehicle open field trials. On day 32, we performed an additional pup exposure with vehicle treatment (Figure 6A). Stressed females had a significant difference in weight compared to control females (Figure 6B). During open field, CNO administration did not induce changes in exploratory behaviors in either group (Figure 6C-F). In the pup exposure assay, both stressed and control virgin females had improved cumulative retrieval with CNO treatment, which we did not observe in the Cre negative group (Figure 6 G, H; Supplemental Figure 6-1). Strikingly, stressed females treated with CNO displayed improved latency to retrieve, time spent crouching, time spent in nest and overall time spent parenting which did not occur in unstressed controls or Cre negative controls, or in the final vehicle session (Figure 6I, L, M, N; Supplemental Figure 6-1). No significant changes were observed in pup grooming (Figure 6K). Interestingly, pup investigation was significantly decreased in both groups which may be due to increased familiarity (Figure 6J) (Bielsky *et al*., 2005; Richter *et al*., 2005; Moy *et al*., 2008). CNO administration did not improve any of the parenting measures in stressed Cre negative females (Supplemental Figure 6-1). We confirmed that CNO administration reduced activity in the PeFA^Ucn3^ neurons using both histology and fiber photometry recording (Supplemental Figure 6-2). Altogether, inhibition of PeFA^Ucn3^ neuronal activity leads to enhancement in parenting behaviors in stressed virgin females (Figure 6O, P).

**Figure 6.**
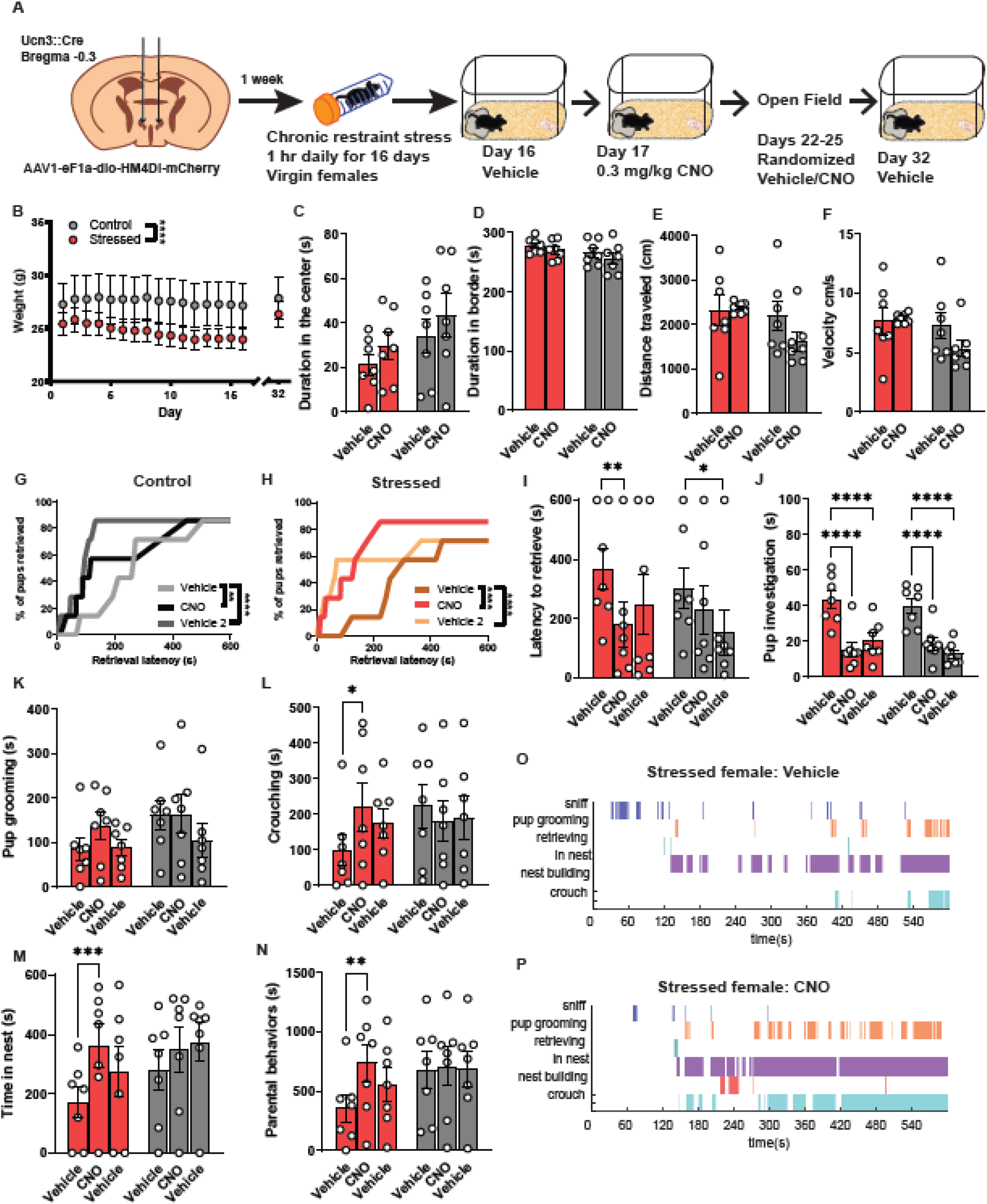
PeFA^Ucn3^ inhibition ameliorates parenting deficits in stressed virgin females. **(A)** Schematic of viral injection strategy and behavior timeline (n=7 stressed Cre+ females; n=7 control Cre+ females). **(B)** Stressed females had a significant difference in weight compared to control females (2-way repeated measures ANOVA, main effect of stress x time p<0.0001). **(C-F)** Open field results show no difference in time spent in center, border, distance moved, or velocity in control or stressed mice with or without CNO treatment. **(G)** Cumulative pup retrieval in control animals improves significantly with CNO injection (Friedman’s test; p=0.0013). **(H)** Cumulative pup retrieval in stressed animals improve with CNO administration (Friedman’s test; p=0.0003). **(I)** CNO treatment decreases latency to retrieve in stressed females (Two-way repeated measures ANOVA, main effect time F_(2,24)_=8.978 p<0.01; Sidak’s multiple comparisons test) but not in control animals. **(J)** Pup investigation dramatically reduces in both animal groups with CNO (Two-way repeated measures ANOVA, main effect of time F_(2,24)_=82.86 p<0.0001; Sidak’s multiple comparisons test). **(K)** Pup grooming is unchanged with CNO treatment. **(L)** Time spent crouching increases with CNO administration in stressed females (Two-way repeated measures ANOVA, main interaction effect of stress and drug F_(2,24)_=3.506 p<0.0462_)_; Tukey’s multiple comparisons). **(M)** Stressed females spent significantly more time in the nest with CNO administration (Two-way repeated measures ANOVA, main effect of time F_(2,24)_=9.399 p=0.001; Sidak’s multiple comparison’s test). **(N)** Cumulative time spent parenting increases with CNO treatment in stressed females (Two-way ANOVA, main effect of time F_(2,24)_=3.942 p=0.0331; Sidak’s multiple comparisons). **(O-P)** Representative behavior traces showing induction of more parenting behaviors with CNO in stressed females. (Significant post-hoc comparisons noted as follows: *p=0.05; **p=0.01; ***p=0.001; ****p=0.0001).

## Discussion

Previously, we have observed that virgin females showing alloparental behavior toward pups have a low-level of activation of PeFA^Ucn3^ neurons while activating these cells highly with chemogenetic or optogenetic methods leads to infant-directed neglect and aggression toward pups (Autry *et al*., 2021). In the present study, we further explored the role of urocortin-3 neurons of the perifornical area during female alloparental and maternal behavior. We aimed to examine the intersection between the role of these neurons in infant-directed behavior and their putative role in the hypothalamic-pituitary-adrenal axis. We hypothesized that PeFA^Ucn3^ neurons become more active with stress and increased PeFA^Ucn3^ cell activity would lead to deficits in pup-directed behavior. Therefore, we studied activation levels of PeFA^Ucn3^ neurons in virgin females and mothers exposed to pups with and without stress and studied the effect of PeFA^Ucn3^ neuron inhibition on alloparental behavior under non-stressed and stressed conditions.

We find that around 20% of PeFA^Ucn3^ neurons are active during infant-directed behavior in virgin females, replicating our previous findings (Autry *et al*., 2021). In addition, we replicated our finding that mothers have a lower level of PeFA^Ucn3^ neuronal activation during pup exposure compared to virgin females. However, we noticed that the controls for our mother group, in which we typically remove the litter, had a high baseline of PeFA^Ucn3^ neuron activity compared to virgin females exposed to bedding. We therefore added a group of mothers who did not have their litters removed. This experiment revealed that our control bedding exposure and experimental pup exposure conditions impact PeFA^Ucn3^ neural activity differentially in mothers depending on whether the mother’s litter is present. These results indicate that PeFA^Ucn3^ neurons may be sensitive to social contexts; these neurons appear to have high baseline activity in mothers with their litters removed during a control bedding exposure and this activity level plummets with introduction of foreign pups. In the future, it will be important to tease apart whether this heightened PeFA^Ucn3^ neural activity after litter removal may be related to an aversive or stressed state, or possibly a social motivation set point that is altered after a female gives birth. Indeed, previous studies have shown that maternal separation can impact both a mother and their offspring’s behavior in measures related to anxiety, social behavior, and cognition (Lemaire *et al*., 2000; Weinstock, 2001; Chapillon *et al*., 2002).

We next assessed the impact of inhibiting this low-level of activity during infant-directed behavior. We found that there was a subtle but significant effect of inhibition PeFA^Ucn3^ neurons on pup-retrieval latency in virgin females. We previously observed that activation of the excitatory PeFA^Ucn3^ neuron projections to the ventromedial hypothalamus or lateral septum mediate infant avoidance and neglect. Our current results suggest that inhibition of PeFA^Ucn3^ neurons in virgin females may lead to decreased activity in these target areas responsible for negative pup directed behavior, allowing for faster infant retrieval. While the behavioral impact of this manipulation is relatively minor, it is in line with the low-level of activation we observe at the cellular level.

In a parallel set of experiments, we tested the impact of chronic stress on infant-directed behavior and PeFA^Ucn3^ neuronal activity in virgin females and mothers. We started by testing a chronic restraint stress paradigm in virgin females. With this paradigm, we observed significant weight loss in females exposed to daily restraint compared to unstressed females. Prior to testing infant-directed behavior, we tested open-field behavior to ensure that the stress paradigm had a behavioral impact after two weeks of chronic restraint, and indeed we observed a reduction in exploration time of the center of the arena in stressed females. Thus, we continued with the infant-directed behavior assay and observed that females with chronic restraint showed significantly less pup retrieval compared to unstressed females, with 2 out of 9 stressed females displaying infant-directed aggression behavior. When we examined the physiological effects of stress, we found that circulating CORT levels were not impacted though PVH^CRF^ neuron activation was significantly reduced in females subjected to stress, suggestive of HPA axis habituation to the repeated stress. However, we did find enhanced PeFA^Ucn3^ neuron activity in chronically stressed females. Intriguingly, we found a strong negative correlation between PeFA^Ucn3^ neuronal activity levels and overall alloparenting behavior in unstressed females, and this correlation was lost in stressed females. Together, these data support our hypothesis that stress increases activity of PeFA^Ucn3^ neurons, and heightened activity in PeFA^Ucn3^ neurons negatively impacts infant-directed behavior.

We therefore proceeded with the chronic restraint paradigm in lactating females. We used a similar timeline for testing, with daily weighing and an open field test prior to pup exposure. We observed significant weight loss in lactating females exposed to stress, however our open field test revealed that females with chronic restraint stress displayed increased exploration of the center of the open field relative to control females. This behavior may be a sign of hypervigilance and may be explained by an increase in distance traveled as well as velocity (Cabib *et al*., 1988; Sequeira-Cordero *et al*., 2019; Rudolph *et al*., 2020). In our subsequent infant-directed behavior assessment, we found no differences in maternal behavior between unstressed and chronically restrained mothers. In our physiological measures, we observed that CORT levels were decreased in mothers with chronic restraint stress with a decrease in both CRF and Ucn3 neuronal activation. The decrease in CORT and CRF neuronal activation are indicative of HPA axis habituation to the repeated stress. Indeed, previous studies have illustrated a reduction in CRF/Fos colabeling or electrophysiological properties of CRF neurons in the PVH with repeated restraint stress in rodents (Bonaz & Rivest, 1998; Matovic *et al*., 2020). We interpret the decrease in Ucn3 neuronal activity as a protective mechanism, preserving maternal behavior under stressful conditions. Indeed, previous studies show that lactating mothers display changes in HPA axis responsivity to stress (Johnstone *et al*., 2000; Douglas *et al*., 2003; Klampfl & Bosch, 2019), and reduction in PeFA^Ucn3^ neuronal activity may contribute to behavioral adaptations to stress.

To overcome the habituation to repeated restraint we observed in mothers, we attempted to perform chronic social stress in lactating females (Supplemental Figure 4-1). However, the stress did not result in weight changes or impact maternal behavior. In the future, we hope to identify a stress paradigm for lactating females that impacts maternal behavior. Furthermore, we aim to study the role of PeFA^Ucn3^ neurons in HPA axis hypo-responsivity in mothers to uncover their potential role in maternal behavior preservation.

We plotted female infant-directed behaviors as ethograms to gain broader insight into how stress affects this complex interaction. Surprisingly, we found that virgin females display longer bouts of infant-directed behaviors during the pup exposure assay relative to mothers, whose behavioral motifs appear to be more sporadic from one behavior to the next (Zoubovsky *et al*., 2020). We suspect that this difference is at least in part due to the removal of the litter during habituation for mothers that is not required for testing virgin females. However, this result does imply that, at least in terms of studying the neural architecture of infant-directed behavior, (1) we can collect a rich dataset from virgin females and (2) that we must be diligent in considering the conditions under which we test behavior in mothers in the laboratory given that litter removal may have a significant impact on some experimental parameters (Lonstein, 2005; Smith & Lonstein, 2008; Miller *et al*., 2011).

Finally, we tested if we could rescue stress-induced deficits in alloparental behavior by inhibiting PeFA^Ucn3^ neurons. We confirmed that our chronic restraint stress paradigm led to weight reductions and proceeded with our pup exposure assay. We found that chemogenetic inhibition of PeFA^Ucn3^ neurons did lead to improved alloparental behavior on several measures including latency to retrieve, time in nest, and crouching. We designed the experiment to observe the effect of stress alone on the first day with vehicle administration followed by inhibition of PeFA^Ucn3^ neurons by CNO treatment on the second day, with a final vehicle test after drug washout. We settled on this design based on our previous observation that blocking PeFA^Ucn3^ neuronal activity optogenetically led to prolonged improvement of infant-directed behavior. We included two control groups, the nonstressed Cre positive group, and the stressed Cre negative group to control for the effects of repeated testing. Our data illustrate that inhibiting PeFA^Ucn3^ neurons in stressed females leads to more substantial effects on alloparental behavior compared to either control group.

Overall, we find that PeFA^Ucn3^ neuronal activity is higher in females showing lower levels of positive infant-directed behavior, a trend that can be observed regardless of physiological status. Chronic stress leads to reduced alloparental behavior accompanied by higher numbers of active PeFA^Ucn3^ neurons in virgin females. Blocking activity in PeFA^Ucn3^ neurons rescues infant-directed behavioral deficits in virgin females. Together with previous studies, our results suggest the important role for the level of PeFA^Ucn3^ neuronal activity in the expression of pro- and anti-parental behavior (Supplemental Figure 6-3). These results support the critical role for PeFA^Ucn3^ neurons in the neural circuitry controlling female parental behaviors and the sensitivity of these behaviors to stress.

## Author Contributions

B.A. and A.E.A. designed and performed experiments, analyzed, and plotted data, and interpreted data and wrote the paper. R.A. supported animal experiments and analyzed data. I.C. analyzed and plotted data.

## Acknowledgements and funding

A.E.A. was supported by a NARSAD Young Investigator Award and a Pathway to Independence Award (NIH R00HD085188). B.A. was supported by a diversity supplement to A.E.A.’s NIH award (R00HD085188-S1) and a Tishman Scholarship. We thank Catherine Dulac for intellectual input and project support at Harvard University. We thank Stacey Sullivan for assistance with transfer of data and mice from Harvard University to Albert Einstein College of Medicine. We thank Krysten Garcia for histology assistance. We thank Giovanni Podda for guidance on the photometry analysis. We thank Kostantin Dobrenis, Vladimir Mudragel, and Mariah Marrero for guidance on Axioscan usage. We thank Kevin Fisher for assistance with image export and analysis. We thank all the members of the Autry lab for input on manuscript preparation.

---

**Supplemental Figure 1-1.**
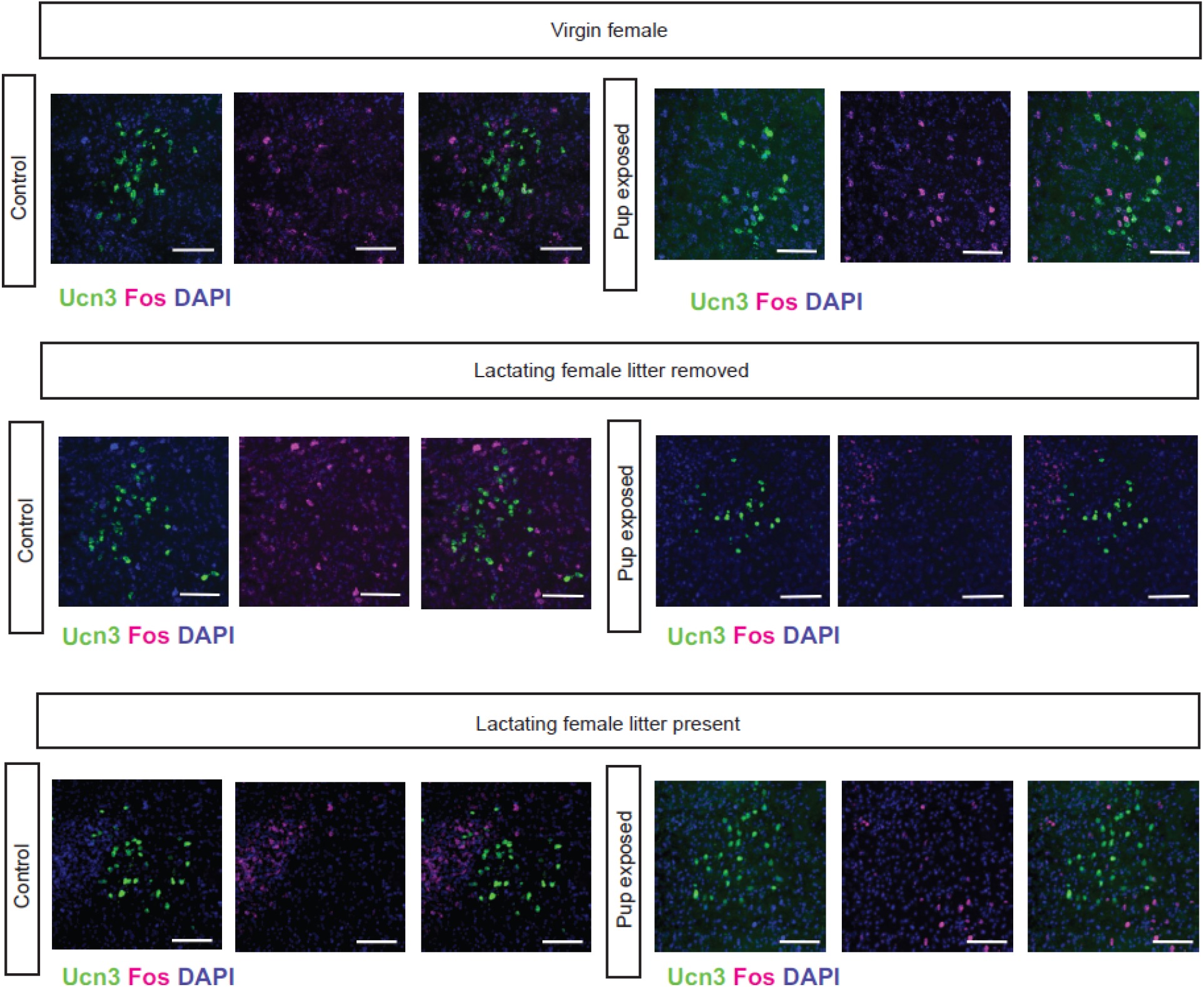
Colocalization of PeFA Urocortin-3 neurons with Fos in response to foreign pups. Representative images from Figure 1 with Ucn3 and Fos channels separated (scale bar 100 µm).

**Supplemental Figure 2-1.**
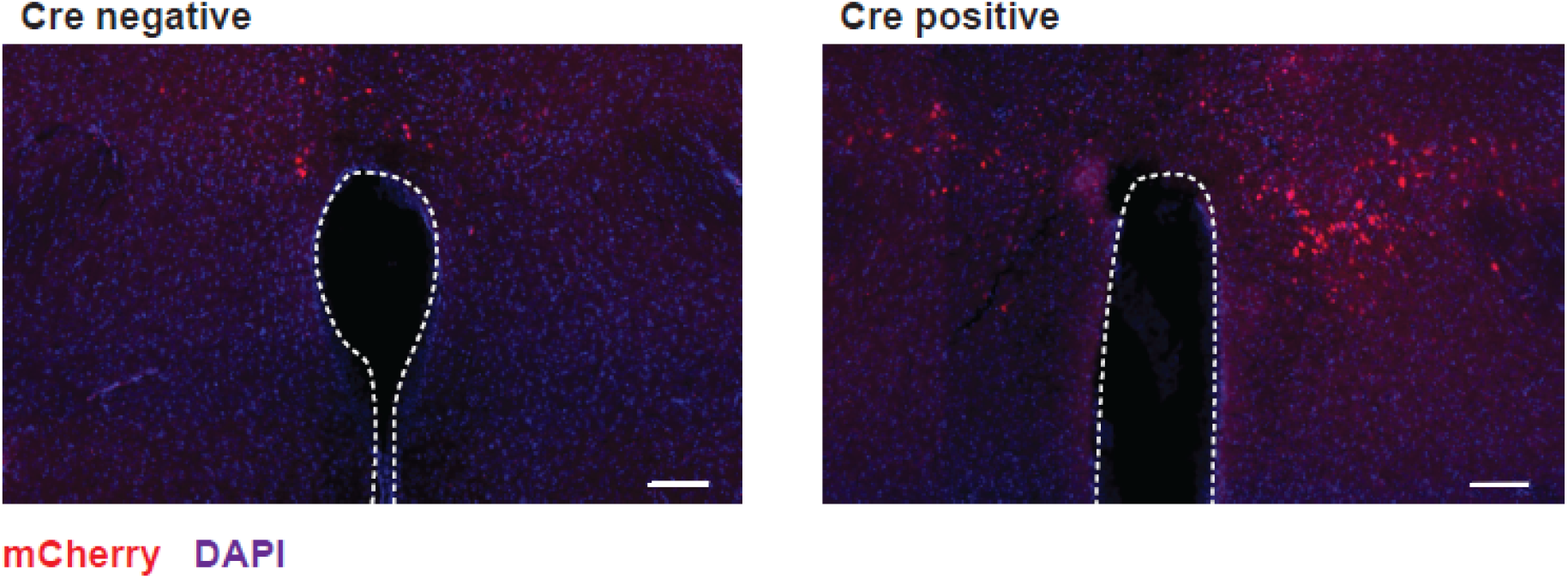
Inhibitory DREADD expression in virgin female mice. Representative image of AAV-mediated hM4di DREADD expression in the perifornical area (PeFA) of Ucn::Cre-(left) or Ucn3::Cre+ (right) females. Third ventricle indicated by outline (scale bar 100 µm).

**Supplemental Figure 4-1.**
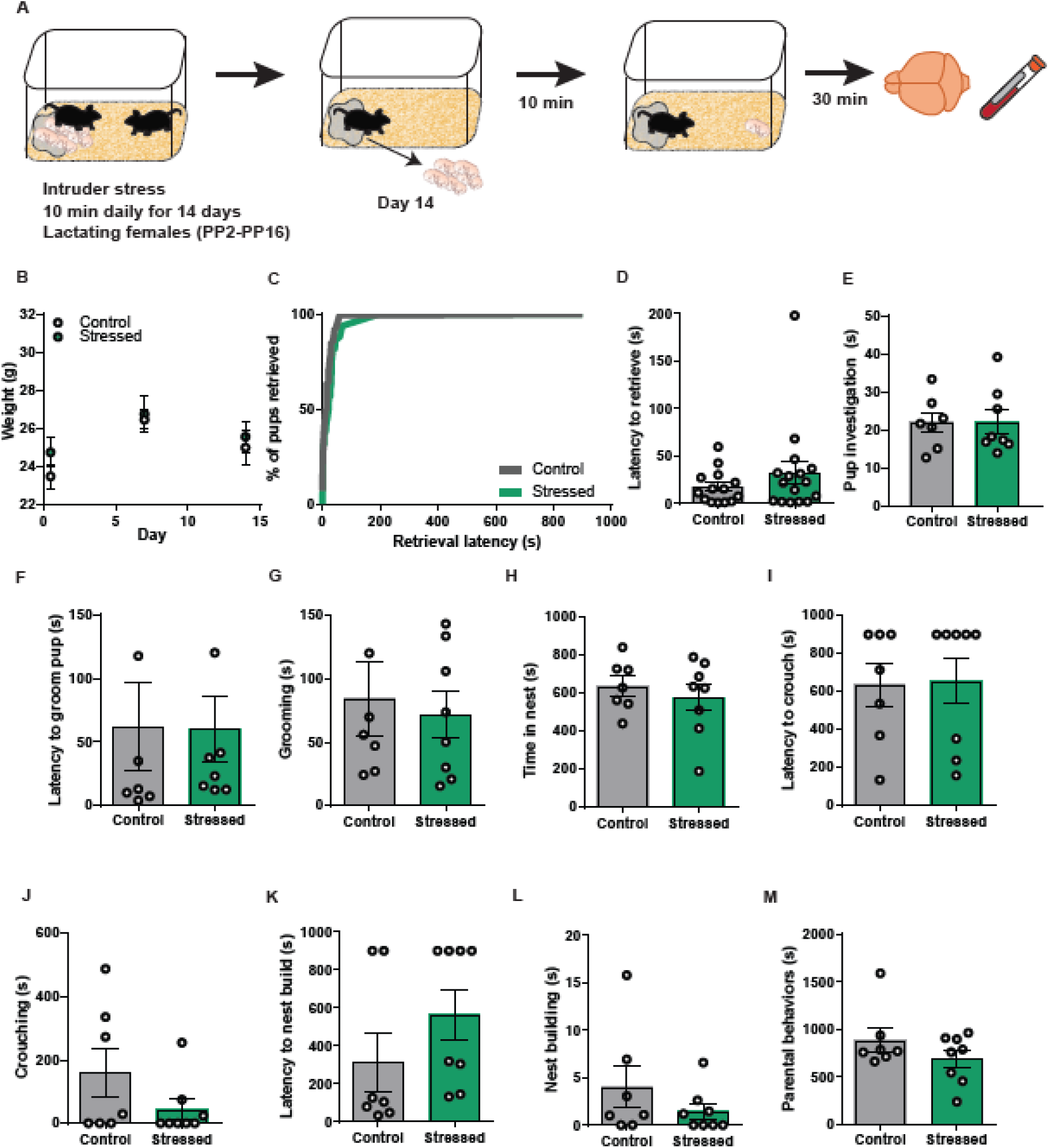
Intruder stress in lactating females does not impact weight or parental behavior. **(A)** Experimental paradigm for intruder stress in lactating females. **(B)** Weight was not affected by stress condition. **(C)** Percentage of pups retrieved was not significantly different between control and stressed females. **(D)** Latency to retrieve **(E)** Pup investigation **(F)** Latency to groom **(G)** Grooming duration **(H)** Time in the nest **(I)** Latency to crouch **(J)** Crouching duration **(K)** Latency to nest build **(L)** Nest building duration and **(M)** Parental behaviors were unchanged between stress and control mothers.

**Supplemental Figure 5-1.**
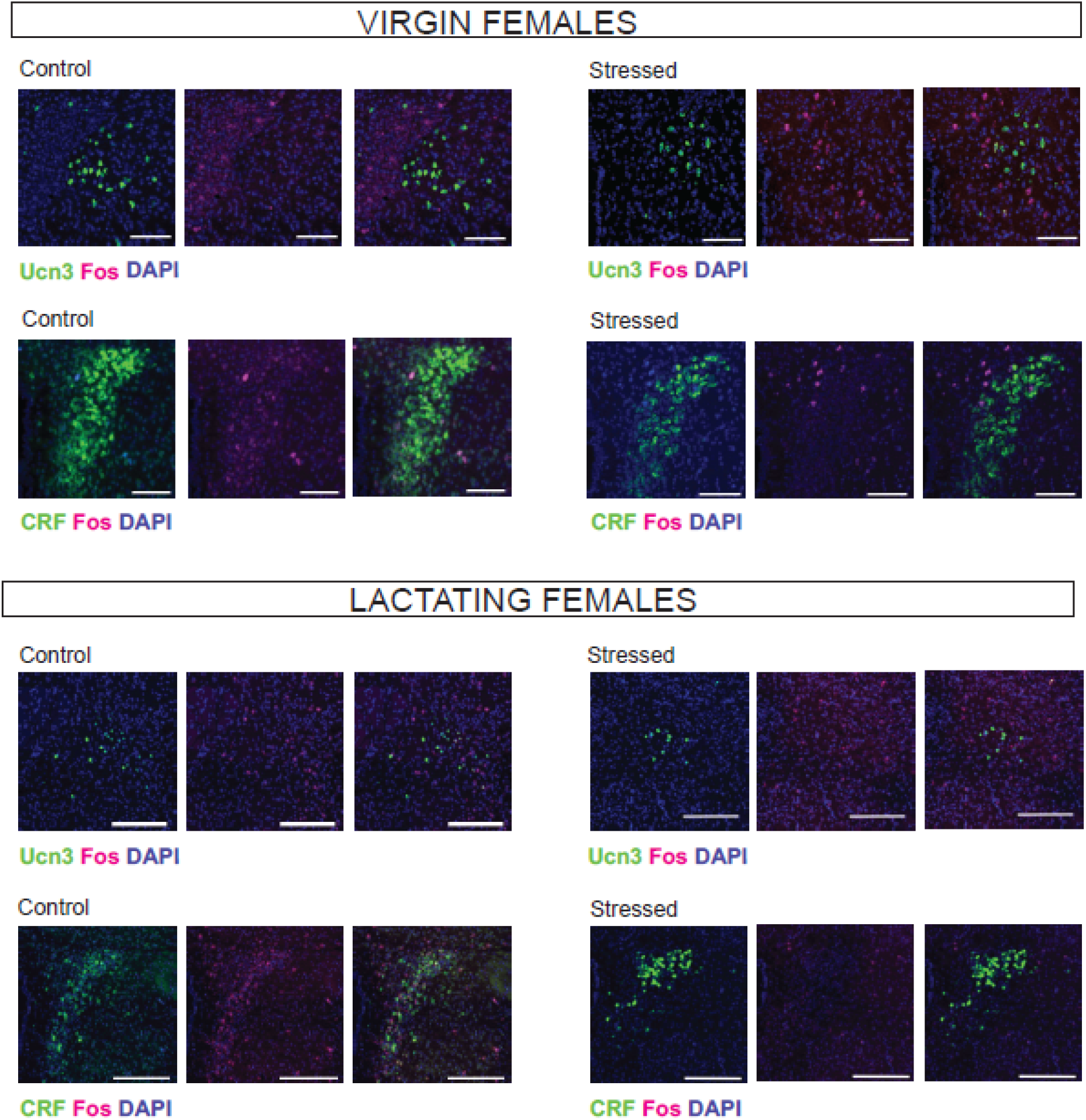
Colocalization of Ucn3 or CRF and Fos in virgin or lactating females exposed to pups in control or chronic restraint stress conditions. Representative images from Figure 5 with Ucn3 or CRF and Fos channels separated (Ucn3 or CRF green channel; Fos magenta channel; scale bar 100 µm).

**Supplemental Figure 5-2.**
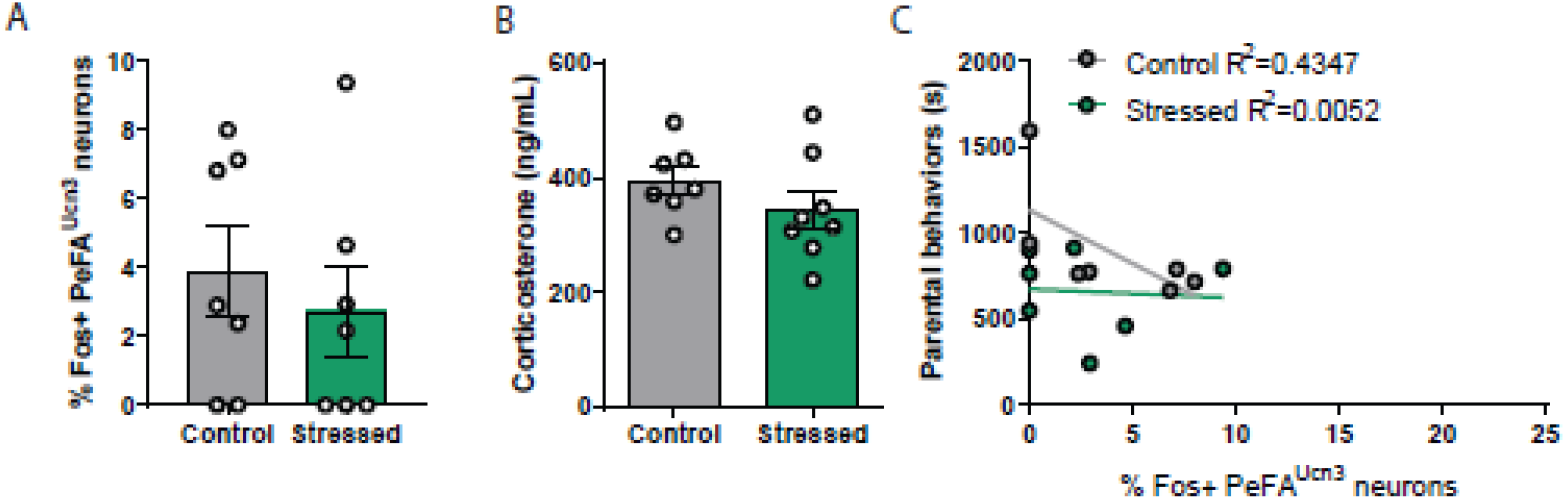
Intruder stress in lactating females does not affect PeFA^Ucn3^ neuronal activation levels or circulating corticosterone. **(A)** Percentage of Fos+ PeFA^Ucn3^ neurons is not significantly different between mothers with or without intruder stress. **(B)** Corticosterone levels are indistinguishable between mothers with or without intruder stress. **(C)** Linear regression analysis reveals a trend towards negative correlation between time spent parenting and PeFA^Ucn3^ activation levels in control lactating females but not in mothers with intruder stress (Control R^2^=0.4347 Stressed R^2^=0.0052).

**Supplemental Figure 6-1.**
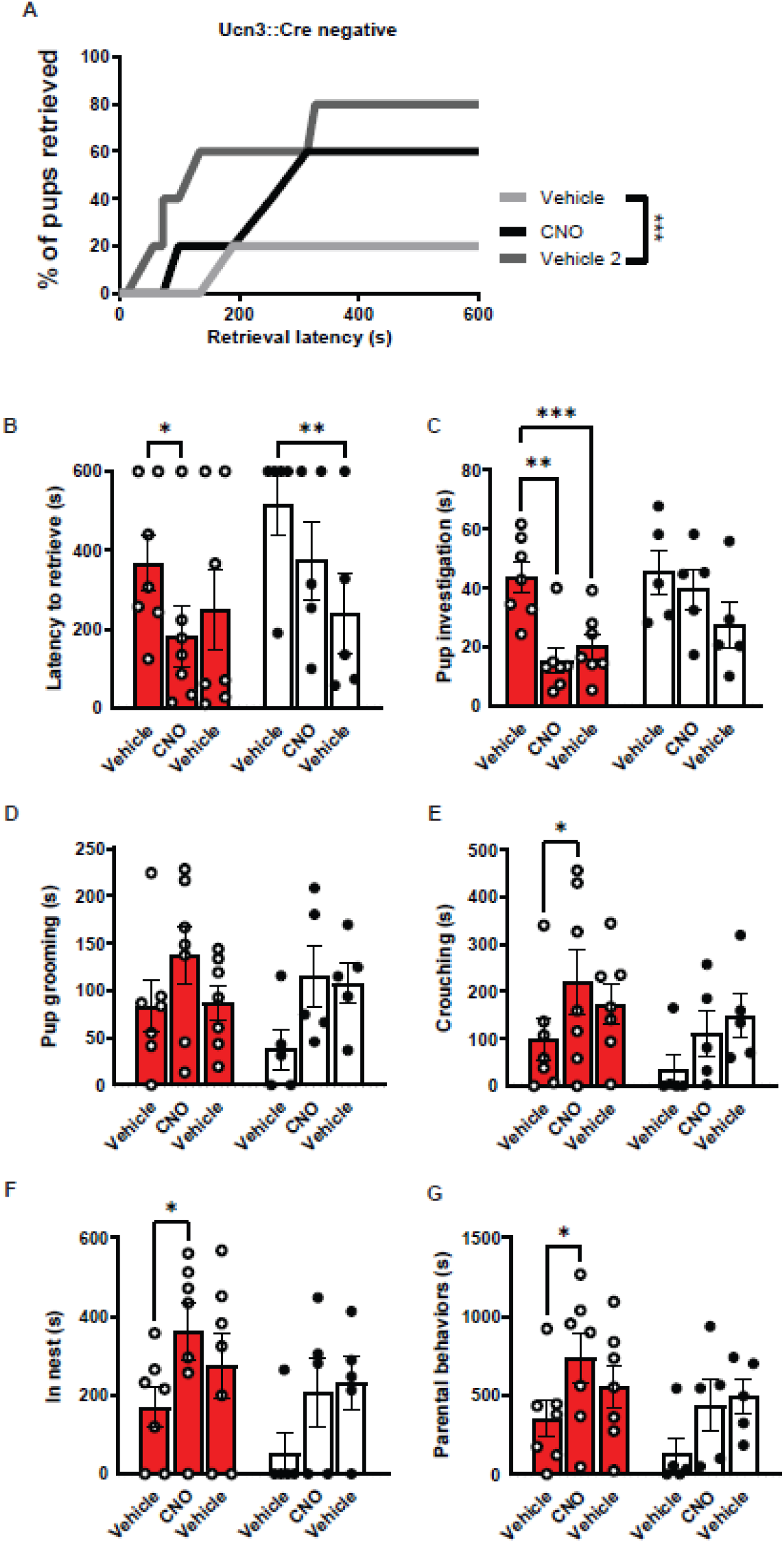
CNO treatment does not impact parenting behavior in Ucn3::Cre-stressed virgin females. **(A)** Cumulative pup retrieval in control animals improves significantly with CNO injection (Friedman’s test; p=0.0002 followed by Dunn’s posthoc comparisons). **(B)** CNO reduces latency to retrieve pups in stressed Ucn3::Cre+ females but not in stressed Cre-females (Two-way repeated measures ANOVA main effect of drug treatment F_(2,20)_=9.667 p<0.0012, Sidak multiple comparisons post-hoc effect significant for Cre+ stressed group vehicle versus CNO p<0.0224 and Cre-stressed group vehicle vs. vehicle 2 p<0.0036). **(C)** Pup investigation is impacted in the stressed Cre+ group but not the stressed Cre-group (Two-way repeated measures ANOVA main effect of drug treatment F_(2,20)_=10.94 p<0.0006, Sidak multiple comparisons post-hoc effect significant for Cre+ stressed group vehicle versus CNO p<0.0010 and vehicle versus vehicle 2 p<0.0009). **(D)** Pup grooming is unaffected. **(E)** CNO treatment increases crouching significantly in Ucn3::Cre positive females (Two-way repeated measures ANOVA main effect of drug treatment F_(2,20)_=4.898, p<0.0186, Tukey multiple comparisons post-hoc effect significant for Cre+ stress group vehicle versus CNO p<0.05) **(F)** as well as duration in nest (Two-way repeated measures ANOVA main effect of drug treatment F_(2,20)_=8.098, p<0.0037, Sidak multiple comparisons post-hoc effect significant for Cre+ stress group vehicle versus CNO p<0.05) **(G)** and cumulative time spent parenting (Two-way repeated measures ANOVA main effect of drug treatment F_(2,20)_=7.728 p<0.0033, Sidak multiple comparisons post-hoc effect significant for Cre+ stress group vehicle versus CNO (p<0.05).

**Supplemental Figure 6-2.**
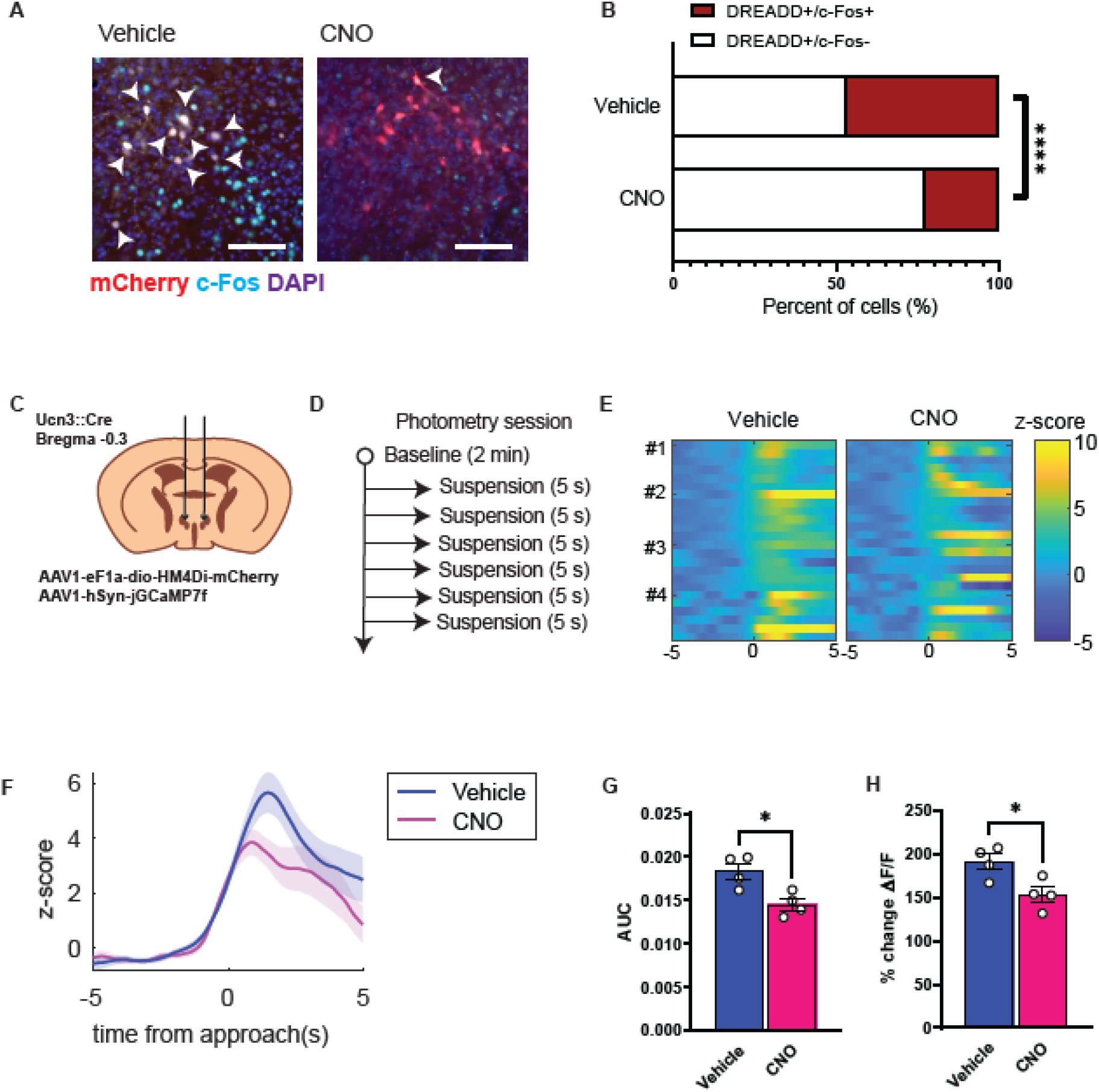
Confirmation of CNO reduction of PeFA^Ucn3^ on AAV-mediated hM4di in Ucn3::Cre females. **(A)** Representative image of changes in PeFA^Ucn3^/c-Fos levels in AAV-hM4di DREADD injected Ucn3::Cre females when CNO is injected intraperitoneally compared to vehicle after 30 minutes of pup interaction (scale bar 100 µm). Arrow indicates Ucn3/c-Fos colocalization. **(B)** Quantification of DREADD expressing neurons co-expressing c-Fos is significantly reduced in Ucn3::Cre females injected with CNO. (Fisher exact test: Vehicle n=180 N=2; Stressed n=101 N=2; p<0.0001). **(C)** Schematic of experimental design to record photometric signals from the PeFA with expression of inhibitory DREADD hM4Di construct in the PeFA^Ucn3^ neurons. **(D)** Schematic of tail suspension session design during photometry recording. **(E)** Z-score of the normalized fluorescence signal (ΔF/F) for each individual trial during the 10s peri-event window centered around the onset of tail suspension. Left: vehicle; right: CNO treatment (n=6 trials, N= 4 female mice) **(F)** Average Z-score of across the 6 trials for each of the four females (dark line represents the mean and the shaded area represents the SEM). **(G)** Area under the curve calculated on the average the tail suspension events (n=6 trials, N=4 female mice). AUC of the 5s segment following the suspension is significantly reduced in the CNO trial compared to vehicle treatment (Paired t-test p<0.0281, N=4 female mice, average of 6 trials for each). **(H)** Percent change of ΔF/F relative to baseline reveals significant reduction of fluorescence evoked by tail suspension with CNO treatment relative to vehicle treatment (Paired t-test p<0.0025, N=4 female mice, average of 6 trials for each).

**Supplemental Figure 6-3.**
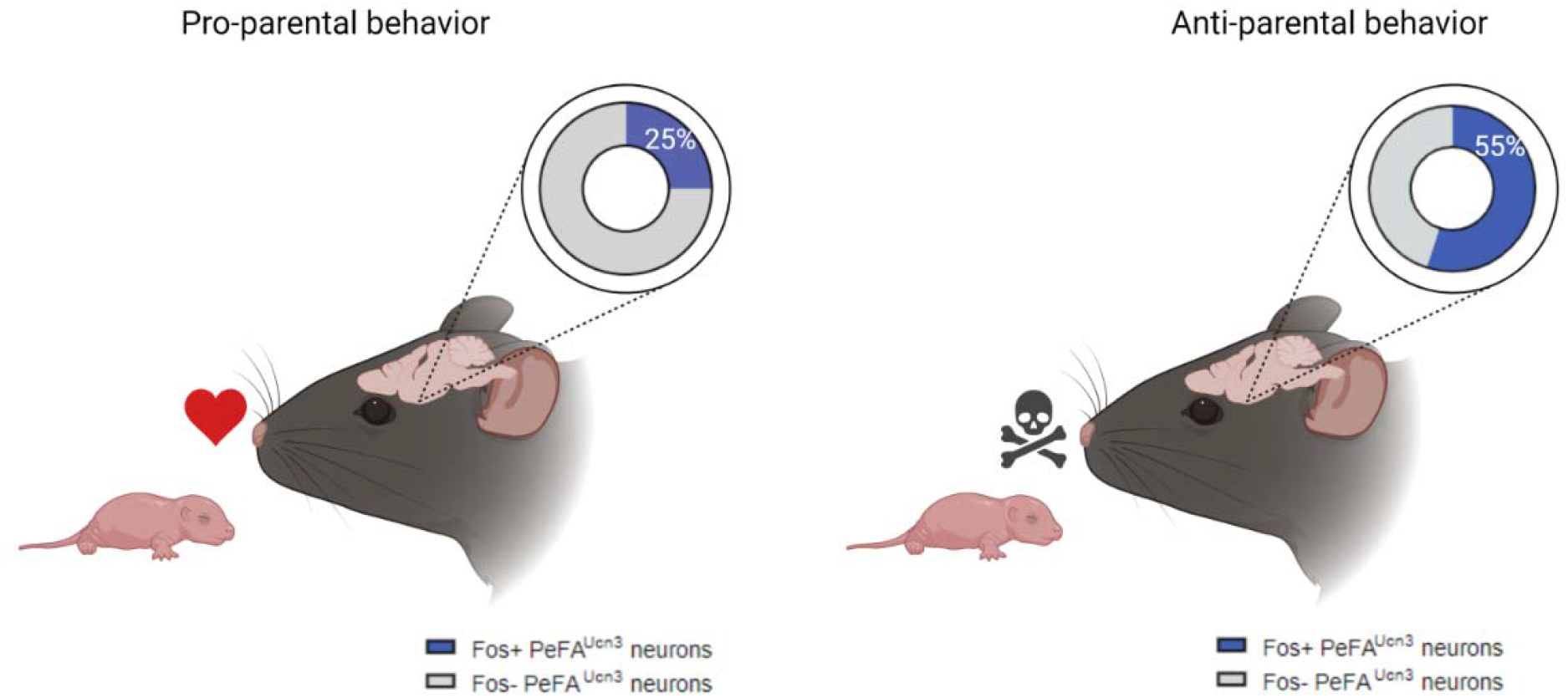
Model for PeFA^Ucn3^ neuronal activation in pup-directed behavior across sexes and physiological states. (Left) Pro-parental behavior such as grooming, retrieving, and crouching over pups in the nest is accompanied by low-level PeFA^Ucn3^ neuronal activation (’25%). Mice in this category include alloparental virgin females, unstressed mothers, and stress-resistant mothers, as demonstrated in this study and Autry et al. 2021. We predict that this level of activation of PeFA^Ucn3^ neurons may also be observed in other categories of mice that display pro-parental behavior including alloparental virgin males, unstressed fathers, and stress-resistant fathers. **(Right)** On the other hand, anti-parental behavior such infant-directed aggression or neglect is accompanied by increased activation of PeFA^Ucn3^ neurons (∼55%). Mice in this category include stressed virgin females and infanticidal males as demonstrated in this study and Autry et al., 2021. We predict that this level of activation of PeFA^Ucn3^ neurons may also be observed in other categories of mice that display anti-parental behavior including stress-susceptible mothers, stressed virgin males, and stress-susceptible fathers. Model created with Biorender.com.

